# Longitudinal Micro-Computed Tomography Detects Onset and Progression of Pulmonary Fibrosis in Conditional *Nedd4-2* Deficient Mice

**DOI:** 10.1101/2023.08.31.555725

**Authors:** Dominik H.W. Leitz, Philip Konietzke, Willi Wagner, Mara Mertiny, Claudia Benke, Thomas Schneider, Wolfram Stiller, Hans-Ulrich Kauczor, Marcus A. Mall, Julia Duerr, Mark O. Wielpütz

## Abstract

**Objectives:** Idiopathic pulmonary fibrosis (IPF) is a fatal lung disease which is usually diagnosed late in advanced stages. Little is known about the subclinical development of IPF. We previously generated a mouse model with conditional *Nedd4-2* deficiency (*Nedd4-2*^−/−^) that develops IPF-like lung disease. The aim of this study was to characterize the onset and progression of IPF-like lung disease in conditional *Nedd4-2*^−/−^ mice by longitudinal micro-computed tomography (CT).

**Methods:** *In vivo* micro-CT was performed longitudinally in control and conditional *Nedd4- 2*^−/−^ mice at 1, 2, 3, 4 and 5 months after doxycycline induction. Further, terminal *in vivo* micro-CT followed by pulmonary function testing and *post mortem* micro-CT was performed in age-matched mice. Micro-CT images were evaluated for pulmonary fibrosis using an adapted fibrosis scoring system.

**Results:** Micro-CT is sensitive to detect onset and progression of pulmonary fibrosis *in vivo* and to quantify distinct radiological IPF-like features along disease development in conditional *Nedd4-2*^−/−^ mice. Nonspecific interstitial alterations were detected from 3 months, whereas key features such as honeycombing-like lesions were detected from 4 months onwards. Pulmonary function inversely correlated with *in vivo* (r=-0.725) and *post mortem* (r=-0.535) micro-CT fibrosis scores.

**Conclusion:** Longitudinal micro-CT enables *in vivo* monitoring of onset and progression and detects radiologic key features of IPF-like lung disease in conditional *Nedd4-2*^−/−^ mice. Our data support micro-CT as sensitive quantitative endpoint for preclinical evaluation of novel antifibrotic strategies.

**NEW & NOTEWORTHY:** IPF diagnosis, particularly in early stages, remains challenging. In this study micro-CT is used in conditional *Nedd4-2*^−/−^ mice to closely monitor the onset and progression of IPF-like lung disease. This allowed us to track for the first time how nonspecific lung lesions develop into key IPF-like features. This approach offers a non-invasive method to monitor pulmonary fibrosis *in vivo*, providing a quantitative endpoint for preclinical evaluation of novel antifibrotic strategies.

## INTRODUCTION

Idiopathic pulmonary fibrosis (IPF) is a fatal chronic progressive lung disease which is characterized by aberrant fibrotic remodeling of the peripheral lung. The resulting progressive loss of lung function combined with limited therapeutic options ultimately leads to respiratory failure (1–5). The diagnosis of IPF is based on the presence of the usual interstitial pneumonia (UIP) pattern in computed tomography (CT) together with the exclusion of known causes of interstitial lung disease using a multidisciplinary diagnostic approach (1). The UIP pattern is characterized by reticular opacities, traction bronchiectasis and subpleural clusters of cystic airspaces described as “honeycombing”, predominantly in the basal and peripheral areas of the lung, which may have varying temporal patterns of occurrence and progression (1, 6). At clinical diagnosis following the onset of symptoms, irreversible lung damage is already present in most patients (7, 8). Therefore, little is known about the onset of IPF and early structural changes have therefore not been defined.

Several mouse models of pulmonary fibrosis exist including bleomycin-induced lung fibrosis (BILF), radiation-induced fibrosis, lung-specific transgenic mice, and models using adenoviral vectors for gene overexpression (9–11). These models have the limitation that pulmonary fibrosis is either transient as the BILF model, or restricted to certain aspects of fibrosis as is the case with transgenic models with overexpression of endogenous fibrogenic mediators or harboring gene mutations associated with familial forms of IPF and animal models of progressive pulmonary fibrosis were lacking (12–14). We and others recently found that the expression of *NEDD4-2* is downregulated in IPF (15–18). Nedd4-2 is an E3 ubiquitin protein ligase involved in posttranscriptional regulation by ubiquitination and targeting for proteasomal degradation of proteins implicated in the pathogenesis of lung disease including surfactant protein C (*Sftpc*), the epithelial sodium channel ENaC and Smad2/3, the intracellular mediators of transforming growth factor β (TGFβ) signaling (16, 19–22). By conditional deletion of *Nedd4-2* (conditional *Nedd4-2*^−/−^), we previously generated a mouse model that develops spontaneous, chronically progressive lung disease that recapitulates key features of IPF in patients such as a distinct fibrotic pattern in histology and micro-computed tomography (micro-CT) including honeycombing-like lesions and fibroblast foci-like fibrotic consolidations, resulting in restrictive lung disease in pulmonary function testing and high pulmonary mortality from three months after induction onwards (16). In our previous work, the characterization of the pulmonary phenotype of conditional *Nedd4-2*^−/−^ mice was limited to advanced stages of lung disease four months after doxycycline induction including *post mortem* micro-CT at approximately 15 µm resolution using a modified scoring system derived from the human situation (16, 23, 24).

The aim of this study was to utilize *in vivo* micro-CT imaging to longitudinally assess the onset and progression of IPF-like lung disease in spontaneously breathing conditional *Nedd4- 2*^−/−^ mice compared to littermate controls. *In vivo* was complemented by *post mortem* imaging in age-matched conditional *Nedd4-2*^−/−^ and control mice. A dedicated micro-CT fibrosis scoring system was employed for semi-quantitative evaluation of key IPF-like features such as honeycombing-like lesions, reticulations, and traction bronchiectasis. In addition, we performed pulmonary function testing to assess the relationship between abnormalities in lung structure and function during the development of IPF-like lung disease in this mouse model. Finally, we assessed the longitudinally scanned mice for radiation-induced lung injury to determine the suitability of micro-CT imaging for *in vivo* monitoring of disease development and response to novel antifibrotic treatment strategies.

## MATERIALS AND METHODS

### Experimental animals

All animal studies were approved by the responsible animal protection authority (Project identification number 35-9185.81/G-45/14, Regierungspräsidium Karlsruhe, Karlsruhe, Germany). Mice for conditional deletion of *Nedd4-2* in lung epithelial cells were generated as previously described (16). In brief, mice carrying *Nedd4-2*^fl/fl^ (25) were intercrossed with *CCSP-rtTA2S-M2* line 38 (*CCSP-rtTA2S-M2*) (26) and LC1 mice (27). All three lines were on a C57BL6/N background. Nedd4-2^fl/fl^ littermates that showed no abnormalities in prior studies served as controls (16). Mice were housed in a specific pathogen-free animal facility and had free access to food and water. For induction of the *Nedd4-2* deletion, mice were exposed to 1 mg/ml doxycycline hydrochloride (Sigma, Darmstadt, Germany) dissolved in a 5% sucrose solution supplied as drinking water in light-protected bottles. Doxycycline solutions were prepared freshly and changed at least every 3 days. To longitudinally study the lung phenotype 4-6 weeks old mice were continuously treated with doxycycline solution for the indicated periods. All mice were included in our study, irrespective of gender, yielding a balanced gender distribution in the conditional *Nedd4-2*^−/−^ groups and littermate controls.

### Pulmonary function testing

Mice were anesthetized using sodium pentobarbital (80 mg/kg), tracheotomized, and placed on the FlexiVent system (SCIREQ, Montreal, QC, Canada). After relaxation with pancuronium bromide (0.5 mg/kg), mice were ventilated with a tidal volume of 8 ml/kg at a frequency of 150 breaths/min and a positive end expiratory pressure of 3 cm H_2_O. The static compliance was derived from pressure–volume curves as described previously (16, 28–33). Mice were then sacrificed for *post mortem* micro-CT.

### Micro-computed tomography

*In vivo* longitudinal micro-CT was performed in conditional *Nedd4-2*^−/−^ mice and littermate controls at 1, 2, 3, 4, and 5 months after induction in supine position using a desktop small animal micro-CT (SkyScan 1176, software version 1.1, Kontich, Belgium). Mice were sedated with 2% isoflurane/2l O _2_ flow inhalation anesthesia via a nasal cone. Scans were acquired in free-breathing with following parameters: 50 kVp X-ray source voltage, 500 mA current, a composite X-ray filter of 0.5 mm aluminum, 55 msec camera exposure time per projection, 720 projections per view, 30 x 30 mm² field of view, acquiring projections with 0.5° increments over a total angle of 360°, resulting in an isotropic pixel size of 35 µm with 1000 x 1000 pixel in plane. The scanning time was approximately 12 min. During image acquisition, the respiratory movements of the thorax were recorded with a visual camera and this information was converted into a pseudosinusoidal signal allowing retrospective respiratory gating. The duration of a complete respiratory cycle was divided into 4 phases of equal length, from the first inspiration to the late expiration. All images were assigned to the corresponding respiratory phase.

After *in vivo* imaging and pulmonary function testing, *post mortem* micro-CT at higher resolution were performed at each timepoint. Mice were sacrificed and again placed in the micro-CT. A fixed pressure to 25 cm H _2_O was applied via the tracheal cannula to ensure sufficient inflation of the lung. Scans were acquired with the following parameters: 50 kVp X-ray source voltage, 550 mA current, a composite X-ray filter of 0.5 mm aluminum, 902 msec camera exposure time per projection, 1440 projections per view, 21 x 31 mm field of view, acquiring projections with 0.25° increments over a total angle of 360°, resulting in an isotropic pixel size of 9 µm with 4000 x 4000 pixel in plane. Total scanning time was approx. 124 min.

Tomograms were reconstructed using NRecon software (version 1.7.0.4, SkyScan, Kontich, Belgium). Reconstruction parameters were smoothing ‘‘2’’, beam-hardening correction ‘‘44%’’; post-alignment and ring artifact correction were manually optimized for each individual scan. For 3D visualization of the distribution of fibrotic lesions the image data sets were processed using VGStudio Max software (version 3.4.3, Volume Graphics GmbH, Heidelberg, Germany). Surface-rendering was computed based on differences in density.

### Micro-CT fibrosis scoring system

Micro-CT was evaluated for pulmonary fibrosis using an adapted fibrosis scoring system as described previously (16, 23, 24) using CTAn software (version 1.10.0.0, SkyScan, Kontich, Belgium) by two independent radiologists (P.K. and T.S.) with 6 and 4 years of experience in thoracic and small animal imaging, respectively. Consensus was achieved by averaging scores from both readers. In brief, lungs were subdivided into 10 equally spaced sections along the longitudinal z-axis form the thoracic inlet to the dome of diaphragm. At each level, lungs were separated into right and left lung according to the perpendicular line that passes the midline of the sternum. For *post mortem* and *in vivo* micro-CT, the scoring system was modified to adapt to the different resolution: The *post mortem* fibrosis score comprised subscores for consolidations, reticular opacities, peripheral bronchiectasis, honeycombing, parenchymal lines and fissural thickening. Consolidation defined as an increased attenuation with obscuration of the pulmonary vascular markings, and reticular opacities defined as a linear interstitial opacity and thickening of the peripheral connective tissue septa, were scored on a scale from 0 to 5 depending on the lung area involved: 0: absent; 1: >0% to 10%, 2: >10% to 20%, 3: >20% to 40%, 4: >40% to 60%, and 5: >60%. Honeycombing-like lesions defined the presence of small cystic airspaces with irregularly thickened walls adjacent to the pleural surface, and peripheral bronchiectasis defined as bronchial dilatations with a maximal distance of 1 mm to the pleura were scored on a scale from 0 to 3 depending on the pleural surface involved: 0: absent, 1: >0% to 10%, 2: >10% to 20%, and 3: >20%). Parenchymal lines defined as nonvascular linear structures arising from the pleural surface and fissural thickening were scored on a scale from 0 to 1: 0: absent, and 1: present. The maximum score for 9 µm resolution images was ((10x(2x5) + 10x(2x3) + 10x(2x1)) = 180 for each side and 360 for the total lung.

*In vivo* micro-CT was scored for area of consolidations as described above for the *post mortem* fibrosis score, as well as for the number of consolidations from 0 (absent) to 5, resulting in a maximum score of 10 x (2x5) = 100 for both the right or the left lung, and a maximum score of 200 for the total lung.

### Interobserver reliability

To establish a micro-CT based scoring system for standardized quantification of pulmonary fibrosis with focus on IPF characteristics in mice, we modified a fibrosis score already used in previous publications (23, 24) and adapted it for use in *in vivo* and *post mortem* micro-CT. All images were scored independently by 2 experienced thoracic radiologists and the weighted Kohens kappa (11) was calculated to evaluate interrater reliability (Supplemental Table S1). For *in vivo* micro-CT almost perfect agreement was achieved for numbers and areas of consolidations and a substantial agreement for the *in vivo* total micro-CT fibrosis score. For *post mortem* micro-CT almost perfect interobserver agreement was achieved for consolidations and fissural thickening. The agreement for the total *post mortem* micro-CT fibrosis score was fair (Supplemental Table S1).

### Radiometry

Radiation dose was estimated using a LiF: Mg/Cu/P-thermoluminescent dosimeter (TLD) also called GR-200A (34), individually calibrated with respect to a CS 137 radioactive source of known activity. The TLD was scanned 5 times using the *in vivo* protocol and the registered dose was averaged. The cumulative equivalent dose measured was 1983 mSv *in aqua* after 5 scans, resulting in 393 mSv as an approximation to the dose applied per scan in mice.

### Statistical analyses

All data are shown as mean ± SEM. Data were analyzed with GraphPad Prism version 9 (GraphPad Software Inc, LaJolla, CA, USA). For comparison of two groups, Mann-Whitney test was used. Comparison of more than two groups was performed with Kruskal-Wallis test followed by Dunn’s post hoc test. For multiple comparisons, P values were adjusted according to the Bonferroni method or by using the False Discovery Rate (FDR) method proposed by Benjamini, Krieger, and Yekutieli as appropriate (35, 36). The Spearman rank order correlation coefficient (r) was calculated for micro-CT fibrosis scores and the pulmonary compliance. We used IBM SPSS Statistics version 27.0.1.0 (IBM Corp, Armonk, NY, USA) to calculate the weighted Cohen’s kappa with a squared weight matrix, providing a more accurate assessment of inter-rater reliability for ordinal data with multiple categories.

## RESULTS

### Longitudinal *in vivo* micro-CT imaging detects onset and progression of pulmonary fibrosis in conditional *Nedd4-2*^−/−^ mice

To determine the onset and progression of IPF-like lung disease, we performed longitudinal studies using monthly *in vivo* micro-CT in a cohort of conditional *Nedd4-2*^−/−^ mice and controls (Figure 1a,b). Standardized scoring showed that the fibrosis score was significantly increased at 3 months after doxycycline induction in conditional *Nedd4-2*^−/−^ mice compared with control mice (Figure 1b,c). Micro-CT at 4 months and 5 months showed progressive pulmonary fibrosis with continuously increasing fibrosis scores in conditional *Nedd4-2*^−/−^ mice (Figure 1b,c). 3D reconstructions revealed the spatial distribution of fibrotic lesions in the lungs, which originated in basolateral regions, increased both in number and volume, and extended apically over time. Distinct fibrotic lesions partially merged into massive fibrotic conglomerates (Figure 1d).

**Figure 1.**
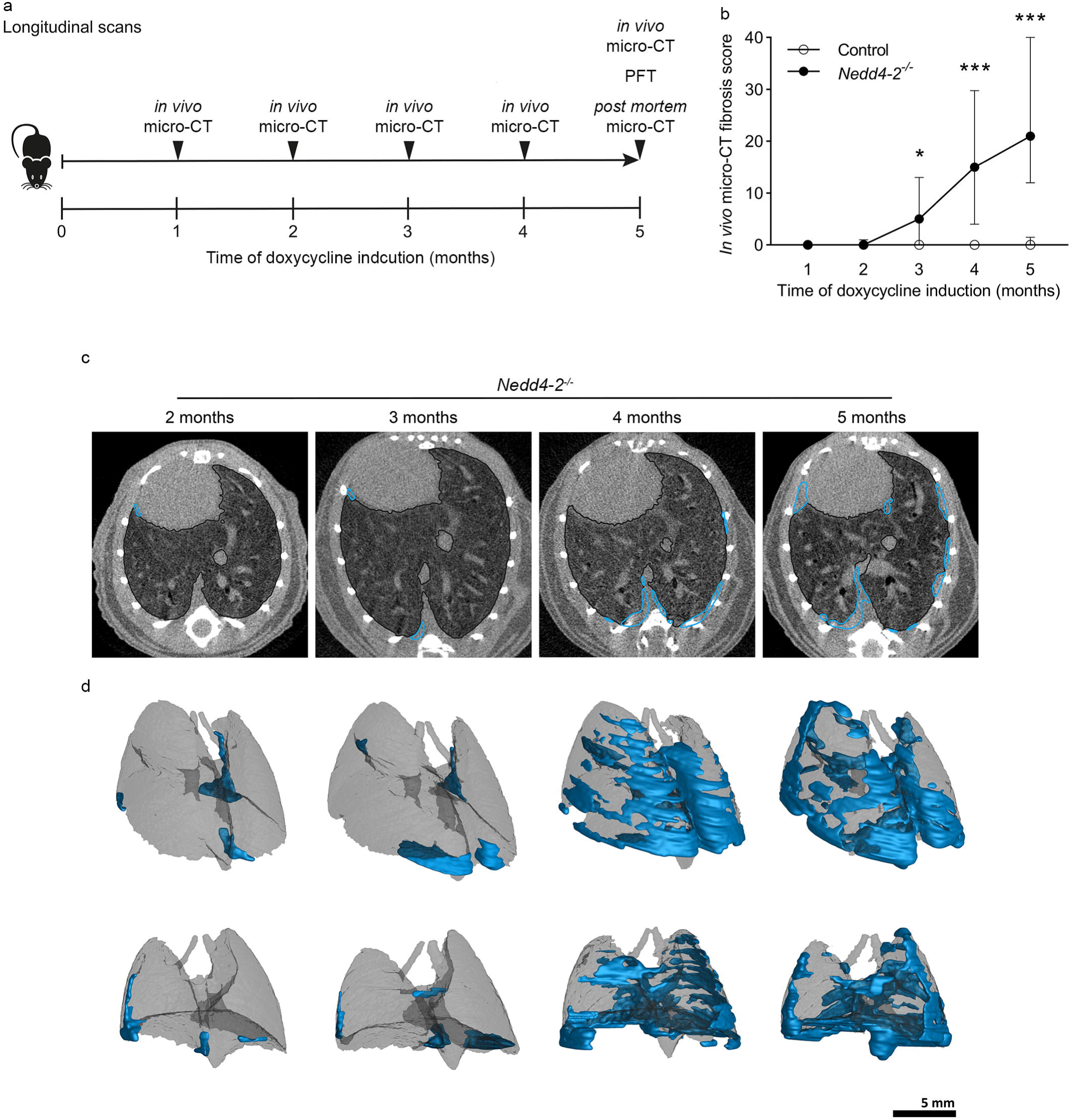
Longitudinal in vivo micro-CT detects onset and progression of pulmonary fibrosis in conditional Nedd4-2^−/−^ mice. (a) Schematic illustration of the study design with serial *in vivo* micro-CT 1, 2, 3, 4 and 5 months after doxycycline induction of conditional *Nedd4-2*^−/−^ mice and controls. (b) Summary of *in vivo* micro-CT fibrosis scores. (c) Representative *in vivo* micro-CT image of a spontaneously breathing conditional *Nedd4-2*^−/−^ mouse scanned at the indicated time points after doxycycline induction. Fibrotic areas are indicated in blue. (d) 3D reconstructions of fibrotic lesions (blue) in the lung of the same mouse as shown in c after 2, 3, 4 and 5 months of doxycycline induction. n = 4-14 mice/group. *\*P < 0.05, ***P < 0.001*.

### Comparison of *in vivo* and *post mortem* micro-CT during the development of IPF-like lung disease in conditional *Nedd4-2*^−/−^ mice

To study the onset and progression of IPF-like lung disease in conditional *Nedd4-2*^−/−^ mice at higher resolution, we performed *in vivo* and *post mortem* micro-CT imaging of conditional *Nedd4-2*^−/−^ mice at the same time points as for the longitudinal *in vivo* studies (n = 5-9 mice per time point) (Figure 2a).

**Figure 2.**
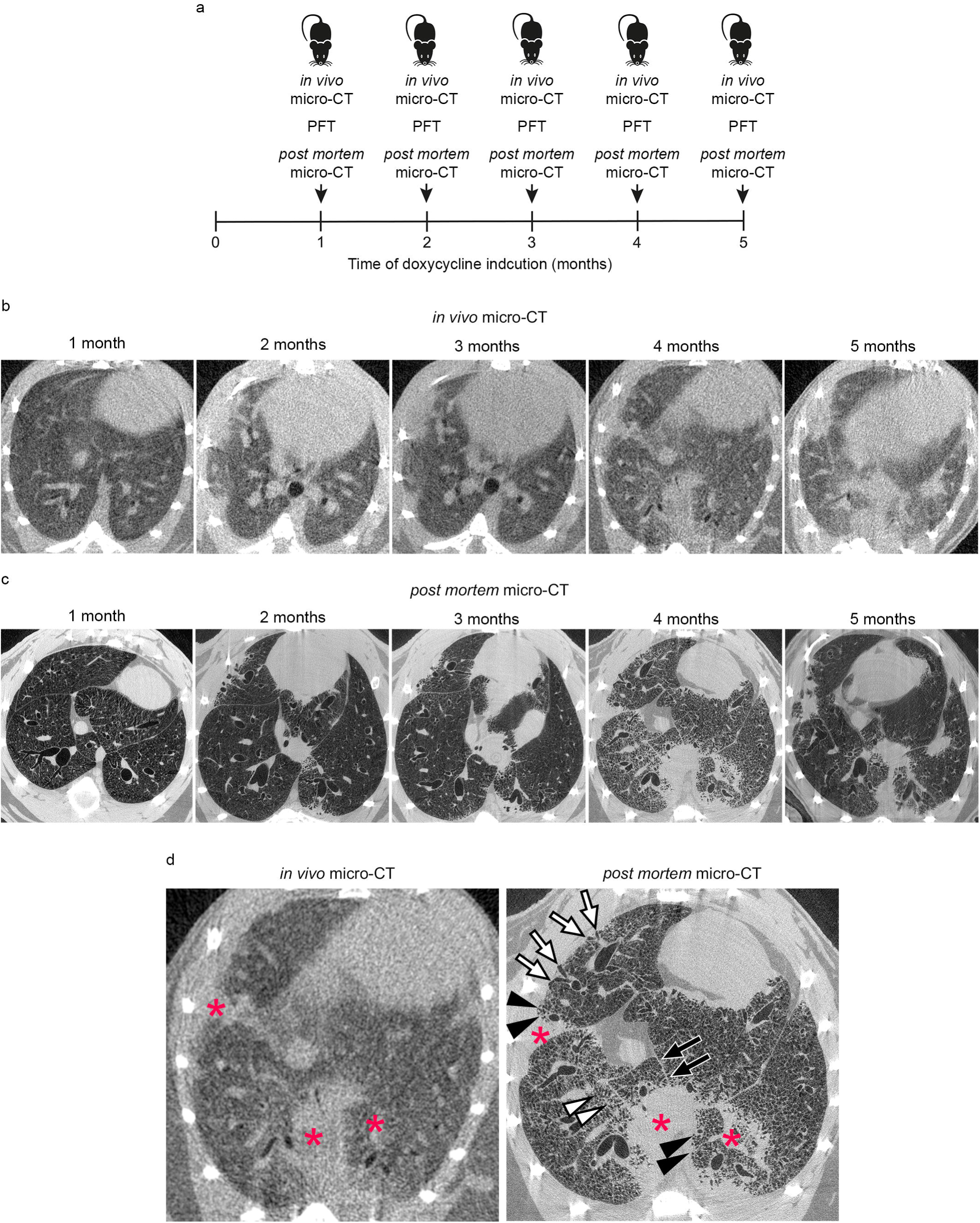
Comparison of radiologic features determined by in vivo and post mortem micro- CT during the development of IPF-like lung disease in conditional Nedd4-2^−/−^ mice. (a) Study design of cross-sectional micro-CT imaging during the development of IPF-like lung disease in conditional *Nedd4-2*^−/−^ mice using *in vivo* micro-CT imaging followed by pulmonary function testing (PFT) and a *post mortem* micro-CT scan after 1, 2, 3, 4 and 5 months of doxycycline induction. (b,c) Representative *in vivo* micro-CT images in spontaneously breathing conditional *Nedd4-2*^−/−^ mice (b) and *post mortem* micro-CT (c) at indicated time points. (d) Comparison of consolidations (red asterisks), honeycombing-like lesions (black arrowheads), peripheral bronchiectasis (white arrows), parenchymal lines (black arrows) and reticulations (white arrowheads) in *in vivo* and *post mortem* micro-CT 4 months after conditional *Nedd4-2* deletion.

Similar to *in vivo* scans, *post mortem* micro-CT imaging detected initial consolidations after 3 months of induction in conditional *Nedd4-2*^−/−^ mice, but not in littermate controls (Figure 2b, c). In addition, characteristic IPF-like structural alterations of the lung such as honeycombing-like lesions, fissural thickening, peripheral bronchiectasis, parenchymal lines and reticulations were distinguishable on *post mortem*, but not o n*in vivo* micro-CT images (Figure 2d). At the higher resolution of *post mortem* micro-CT, we observed that a subset of conditional *Nedd4-2*^−/−^ mice already showed an increased fibrosis score after 2 months of doxycycline induction, and that the fibrosis score was significantly increased from 3 months onwards compared with control mice (Figure 3b).

**Figure 3.**
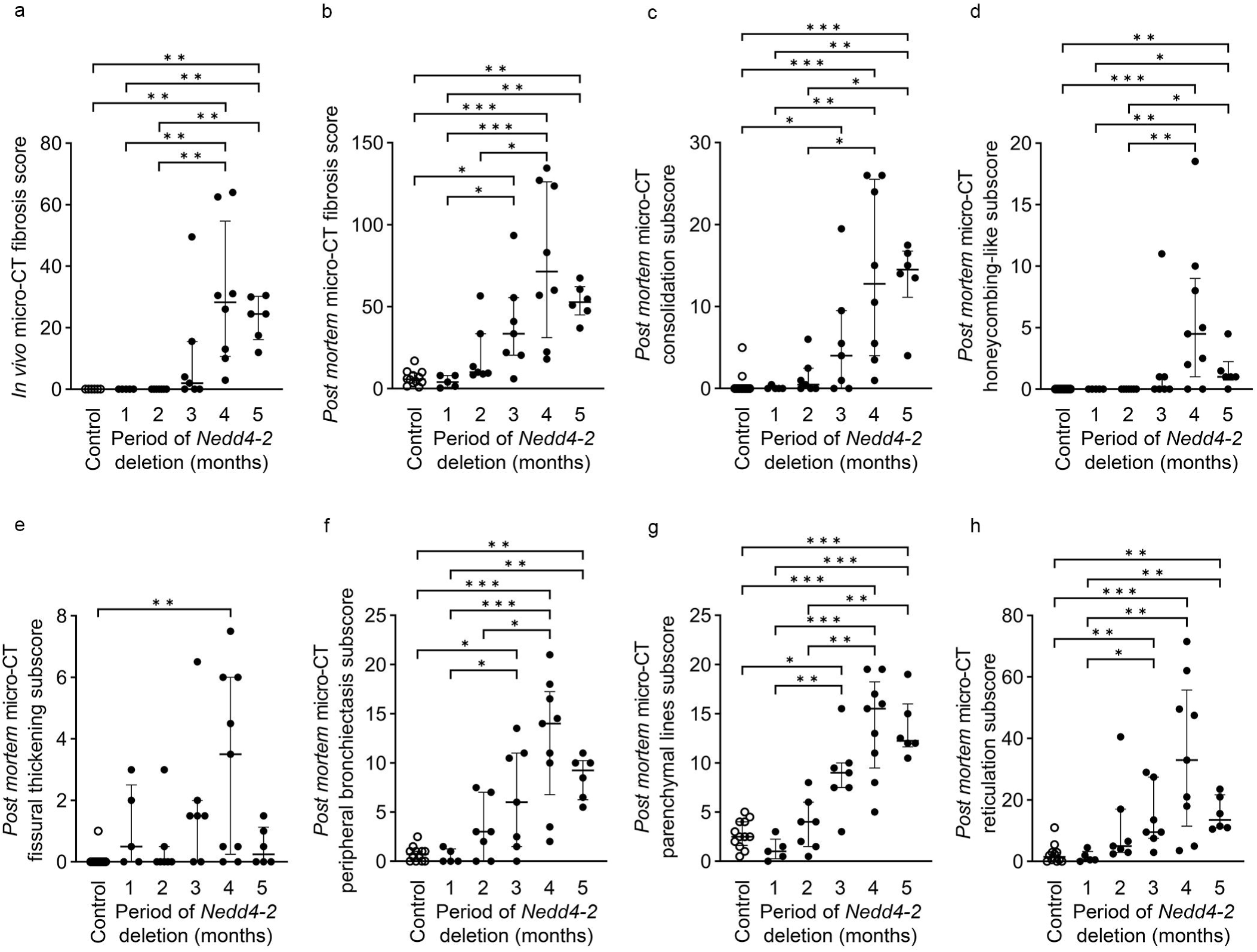
Development of radiologic features of IPF-like lung disease in conditional Nedd4-2^−/−^ mice. (a) *In vivo* micro-CT fibrosis score along disease development after 1, 2, 3, 4 and 5 months of doxycycline induction in conditional *Nedd4-2*^−/−^ mice and controls. (b-h) *Post mortem* micro-CT fibrosis score (b), consolidations (c), honeycombing-like lesions (d), fissural thickening (e), peripheral bronchiectasis (f), parenchymal lines (g) and reticulations (h) detected by high resolution micro-CT in conditional *Nedd4-2*^−/−^ mice and controls. n = 5-12 mice/group. FDR-adjusted *P* values are indicated as *\*P < 0.05, **P < 0.01, ***P < 0.001*.

### Higher resolution *Post mortem* micro-CT detects onset and progression of specific fibrotic lesions in IPF-like lung disease in conditional *Nedd4-2*^−/−^ mice

The higher resolution of *post mortem* micro-CT allowed us to further characterize the onset and progression of fibrotic lesions and distinguish consolidations, honeycombing-like lesions, fissural thickening, peripheral bronchiectasis, parenchymal lines, and reticular structures (Figure 2c and Figure 3b-h). In this cohort, there were no fibrotic changes detectable by in vivo micro CT fibrosis scores in conditional *Nedd4-2*^−/−^ after 1 and 2 months of induction which then increased and achieved significance after 4 and 5 months of induction compared to control mice (Figure 3a). The same mice showed higher post mortem fibrosis scores resulting in a significant difference between conditional *Nedd4-2*^−/−^ and control mice already after 3 months of induction (Figure 3b). The post mortem micro-CT fibrosis subscores revealed a similar dynamic change of consolidations as the in vivo micro-CT fibrosis scores (Figure 3c). These analyses showed that peripheral bronchiectases were already detectable after 2 months in a subset of conditional *Nedd4-2*^−/−^ mice and in all conditional *Nedd4-2*^−/−^ mice after 4 and 5 months of doxycycline induction (Figure 3f). Parenchymal lines and reticulations showed a similar dynamic of which the latter accounted for the largest portion of the total fibrosis score across all age groups (Figure 3b, g, h). Honeycombing-like lesions were mostly seen at later stages of the disease or in very sick mice, and were not observed at 1 and 2 months (Figure 3d). The subscores for fissural thickening were variable along the age groups 1, 2 and 3 months in conditional Nedd4-2 mice and significantly elevated at 4 months in conditional *Nedd4-2*^−/−^ mice compared to control (Figure 3e).

### Micro-CT fibrosis score correlates with lung function impairment in conditional *Nedd4-2*^−/−^ **mice**

Although the *in vivo* micro-CT images did not enable identification of specific fibrotic lesions, the fibrosis scores obtained with *in vivo* and *post mortem* micro-CT images from conditional *Nedd4-2*^−/−^ and control mice showed a strong correlation with each other (r = 0.860, *P* < 0.001) (Figure 4a). To test the relationship between the micro-CT fibrosis score and pulmonary function, we assessed lung compliance prior to *post mortem* micro-C T . Pulmonary function testing demonstrated a significant decrease in lung compliance in conditional *Nedd4-2*^−/−^ mice after 4 and 5 months of induction compared to control mice (Figure 4b) and showed a strong inverse correlation with the fibrosis scores from conditional *Nedd4-2*^−/−^ and control mice obtained from *in vivo* (r = −0.725, *P* < 0.001) and *post mortem* micro-CT images (r = −0.570, *P* < 0.001) (Figure 3b, c).

**Figure 4.**
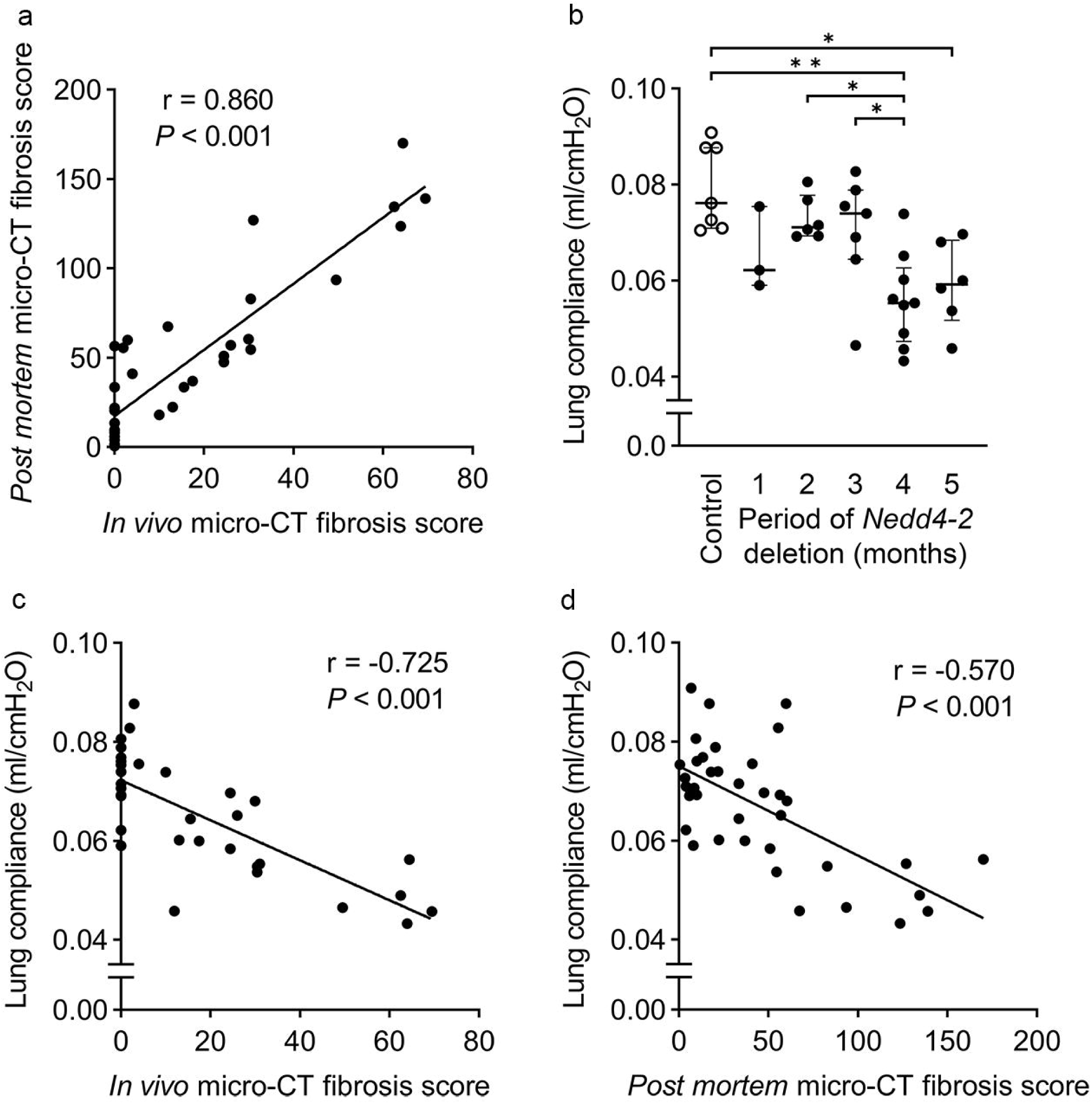
Micro-CT fibrosis score negatively correlates with impairment in pulmonary function in conditional Nedd4-2^−/−^ mice. (a) Correlation between fibrosis scores determined by *in vivo* micro-CT and *post mortem* micro-CT. (b) Summary of lung compliance at different time points in conditional *Nedd4-2*^−/−^ mice compared to controls. Correlation between lung compliance determined by pulmonary function testing and *in vivo* micro-CT fibrosis score (c) and *post mortem* micro-CT fibrosis score (d) in conditional *Nedd4-2*^−/−^ mice. Spearman rank order correlation coefficient (r) and *P*-values are provided. n = 31 mice.

### Micro-CT of end-stage IPF-like lung disease in moribund conditional *Nedd4-2*^−/−^ mice

During our study, 4 conditional *Nedd4-2*^−/−^ mice developed signs of severe respiratory distress with tachypnea and retractions associated with weight loss that had to be euthanized 3 to 4 months after doxycycline induction (Figure 5a). *Post mortem* micro-CT imaging showed that IPF-like lung disease was substantially more severe in this subgroup of moribund mice with end-stage lung disease compared to conditional *Nedd4-2*^−/−^ mice that survived and were studied 5 months post doxycycline induction (Figure 5). Specifically, reticulations and consolidations affected almost the entire lung (Figure 5b) and the *post mortem* micro-CT fibrosis score and all of its subscores were markedly increased in this subgroup of moribund mice compared to conditional *Nedd4-2*^−/−^ mice that were studied at 5 months (Figure 5c-i).

**Figure 5.**
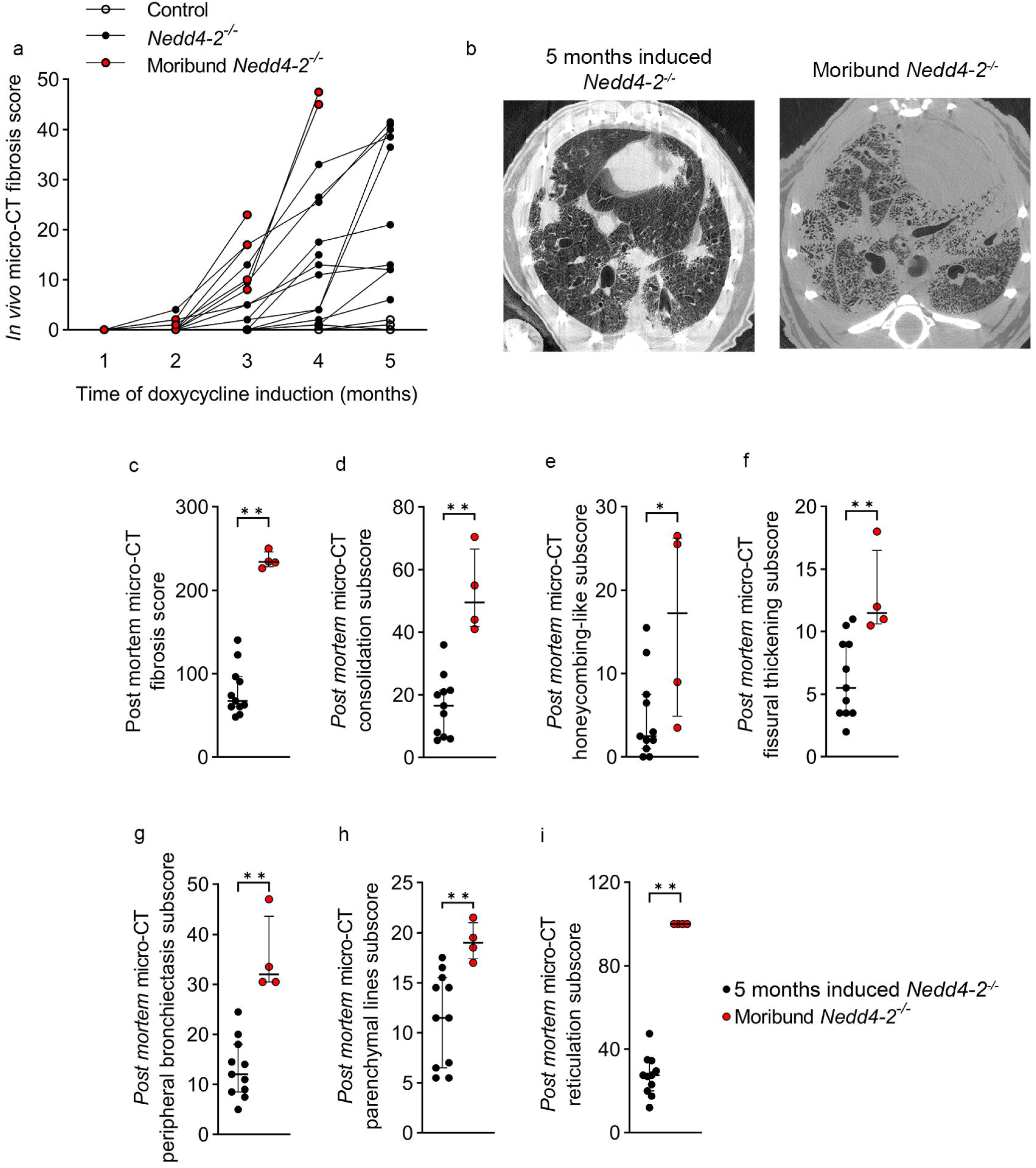
Micro-CT of end-stage IPF-like lung disease in moribund conditional Nedd4-2^−/−^ mice. (a) Development of fibrosis scores in longitudinal micro-CF imaging studies of moribund conditional *Nedd4-2*^−/−^ mice that had to be euthanized vs conditional *Nedd4-2*^−/−^ mice that survived 5 months of doxycycline induction. (b) Representative *post mortem* micro-CT images of a surviving conditional *Nedd4-2*^−/−^ mouse and a moribund mouse that had to be euthanized after 4.5 months. (c-j) Summary of *in vivo* micro-CT fibrosis score (c) the *post mortem* micro-CT fibrosis scores (d) and the subscores for quantification of consolidation (e), honeycombing-like lesions (f), fissural thickening (g), peripheral bronchiectasis (h), parenchymal lines (i) and reticulations (j). n = 5-11 mice/group. *\*P < 0.05, **P < 0.01, ***P < 0.001*.

## DISCUSSION

In this study, we utilized micro-CT imaging to characterize the onset and spontaneous progression of IPF-like lung disease in conditional *Nedd4-2*^−/−^ mice. With longitudinal *in vivo* micro-CT, we were able to closely monitor the development of fibrotic lesions from the onset to end stage lung disease in moribund mice (Figure 1 and Figure 5). Higher resolution *post mortem* micro-CT imaging in age-matched mice provided more detailed insights into the specific structural changes and pattern of disease progression in this model of IPF-like lung disease (Figure 2 and Figure 3). Further, we found that the fibrosis score determined from micro-CT imaging correlates with lung function impairment in conditional *Nedd4-2*^−/−^ mice (Figure 4). Collectively these data suggest micro-CT imaging in conditional *Nedd4-2*^−/−^ mice for non-invasive testing of *in vivo* effects of new therapeutic approaches or disease-relevant factors in early and late IPF-like disease.

This longitudinal study extends our previous micro-CT imaging studies of advanced lung disease in conditional *Nedd4-2*^−/−^ mice and provides important insights into the onset and progression of lung disease in this model of IPF (16). To date, there is little knowledge of the initial lesions of IPF and how they develop *in vivo* into the massive destruction of lung tissue that is observed in patients with IPF. We therefore used the conditional *Nedd4-2*^−/−^ mice to obtain insights into the in vivo development of IPF-like lung disease (16). Similar to previous studies in conditional *Nedd4-2*^−/−^ mice with advanced IPF-like lung disease, we found radiomorphological characteristics of consolidations, reticulations, traction bronchiectasis and honeycombing-like lesions resembling a UIP pattern in this mouse model (Figure 2 and 3) (16). We found that fibrosis scoring based on *in vivo* and *post mortem* micro-CT shows a high correlation with each other (Figure 4), indicating that the use of *in vivo* micro - CT imaging is useful to reliably assess disease progression and potentially response to therapy. However, *in vivo* micro-CT allowed only limited information about detailed structural characteristics of fibrotic changes due to motion artifacts caused by cardiac and respiratory motion as well as a somewhat lower resolution of 35 µm. The subsequent *post mortem* micro-CT with a resolution of 9 µm provided a more detailed characterization of the morphological changes in this model of IPF-like lung disease. Regarding the sequence of disease development, rather subtle and nonspecific changes such as peripheral bronchiectasis or mild reticulations are present in some mice already after 2 months of deletion of conditional *Nedd4-2*^−/−^ and progress over time. Honeycombing-like lesions can also be reliably distinguished by micro-CT with advanced IPF-like disease after 4 and 5 months of doxycyline induction (Figure 2, 3 and 5). This observation is consistent with our previous work including CT imaging and histopathology studies at limited time points, in which only mild nonspecific histomorphological changes were detected in two months induced conditional *Nedd4-2*^−/−^ mice, such as mild macrophage-dominated inflammation associated with an increase in septal wall thickening which could only be detected by studies of the ultrastructure of the lung (16, 37). At later time points, after 3 to 4 months of induction, typical histological features of an IPF-like lung disease with honeycombing-like lesions and fibroblastic foci could be observed (16). This suggests that at the beginning of the manifestation of IPF-like lung disease, rather nonspecific, subtle changes determine the morphology, and that the typical UIP pattern already reflects irreversible lung changes of advanced lung disease (Figure 2, 3 and 5).

The comparison of micro-CT scores from mice that received single vs multiple *in vivo* micro-CT scans also allowed us to assess potential effects of radiation-induced lung injury, which is important since it may mimic IPF-like lung changes (38). We attempted to minimize radiation exposure by keeping the scanning time as short as possible and waiving respiratory-triggered examinations as it would require an even higher radiation dose due to longer scan times. We observed increased fissural thickening and reticulation in conditional *Nedd4-2*^−/−^ mice subjected to repeated vs. single micro-CT examinations (Supplemental Fig. S1). Although little is known about the long-term consequences of radiation on IPF, studies of patients with interstitial lung disease who have undergone radiation therapy for malignant lung tumors indicate that they are at higher risk for acute radiation-induced lung injury (39–41). While the subscores of fissural thickening and reticulation showed a similar trend in control animals (Supplemental Fig. S1), our data suggest that the lung tissue of conditional *Nedd4-2*^−/−^ mice may be more susceptible to damaging events than it is the case in healthy lungs. However, the changes noted in the control mice may also be nonspecific post-inflammatory changes because the measured radiation dose of approximately 0.39 Sv per scan for *in vivo* micro-CT is well below previously reported doses which also did not result in significant fibrotic changes (38, 42) .

Importantly our data demonstrate the feasibility of *in vivo* micro-CT imaging without significant radiation-induced lung injury. The strong correlation between lung compliance and micro-CT fibrosis score also reinforces the importance of this method in predicting the functional outcome in mice. In this context, micro-CT offers a distinct advantage over pulmonary function testing, which is invasive in mice and helps to reduce the number of animals needed, in accordance with animal welfare and 3R principles.

In our study, we found striking differences in the pattern and development of pulmonary fibrosis in conditional *Nedd4-2*^−/−^ mice compared to other experimental models of IPF. One of the most commonly used mouse model of BILF (9). Micro-CT studies in these mice showed that fibrotic alterations occured rapidly after bleomycin instillation, occupying primarily the mid-to-upper regions of the lung with a dorsocentric distribution that emanates from bronchovascular areas to the periphery (23, 24, 43–45). We found that the IPF-like lung disease in conditional *Nedd4-2*^−/−^ mice develops spontaneously, progresses over several months in a chronic, initially slowly progressing way that rapidly worsens at the end of disease development. Further fibrotic lesions in these mice exhibit a basolateral distribution originating from the periphery of the lung, thus more closely mirroring the pattern observed in IPF patients (46). Taken together, the conditional *Nedd4-2*^−/−^ mice are distinct from the mouse model of BILF demonstrated by a greater similarity to IPF in patients which has previously also been shown at the histological level with microscopic honeycombing, fibroblast foci-like structures, bronchiolization of peripheral airways and at the molecular level with altered pulmonary expression of Muc5b and a similarly altered proteomic signature as in IPF (16).

Our results also enable a more detailed comparison of radiological features of the development of lung disease in conditional *Nedd4-2*^−/−^ mice and patients with IPF. Radiological features of UIP, the hallmark of IPF, were detailed in the 2018 guidelines for diagnosis of IPF (46) and revised in the new official ATS/ERS/JRS/ALAT clinical practice guideline which opted to retain the four established classifications of CT patterns in IPF: “UIP pattern,” “probably UIP pattern”, “indeterminate for UIP”, “CT findings suggestive of an alternative diagnosis”, which are based on the distribution and features of CT findings (1).

Moreover, apart from IPF and other relatively rare interstitial lung diseases (ILD), incidental interstitial lung abnormalities (ILA) are increasingly reported in cross-sectional imaging studies from smokers, for example in the setting of lung cancer screening (47). Since diagnosis is usually made late, early findings in IPF have not been defined, which also includes a lack of knowledge on actionable findings in the case of asymptomatic and incidentally found ILA at imaging (47, 48). The conditional *Nedd4-2*^−/−^ mice showed all CT features of a UIP pattern (honeycombing with or without traction bronchiectasis and presence of irregular thickening of interlobar septa). The distribution of morphological abnormalities in conditional *Nedd4-2*^−/−^ mice is predominantly subpleural and basal as well as mostly heterogeneous. However, conditional *Nedd4-2*^−/−^ mice also showed consolidations as a substantial feature, which are suggestive of alternative processes, such as consolidating pneumonia. This difference from human IPF could be related to anatomical differences between mice and humans. Since the airways in mice are much smaller, they might be inherently more susceptible to conditions such as atelectasis, which present as consolidations on CT. Notably, since previous work in this mouse model has shown overexpression of Muc5b, particularly in the periphery of the lung, and decreased mucociliary clearance due to increased transepithelial ENaC-mediated Na ^+^ absorption leading to airway surface dehydration, the observed phenomenon could thus represent atelectasis due to impaired mucociliary clearance and/or mucus plugging (16, 49, 50). However, a predominant peribronchovascular pattern with subpleural sparing suggestive for NSIP was not found. Collectively, the micro-CT findings from our study, which highlight predominant UIP features, combined with our previous observations of IPF-signature lesions in histology and shared molecular signatures with IPF patients, support the relevance of conditional *Nedd4-2*^−/−^ mice as a model of IPF, and potentially other ILD and ILA.

In this study, we employed an adapted micro-CT-based fibrosis score for quantifying fibrotic lesions in a standardized manner (23, 24). The decision to use a semi-quantitative visual score was driven by its simplicity and its reliance on established radiologic CT variables (51). However, scoring systems such as this carry the risk of information loss, as they only capture features that meet predefined criteria. In a longitudinal study in which most diagnostic criteria were derived from advanced disease stages with irreversible changes, there is a possibility that precursor lesions may not be adequately represented in the scoring system. This could result in missing crucial clinical insights into disease evolution. While recent advancements have produced computer tools that promise greater precision in detecting lung damage, they also have potential drawbacks, particularly in pulmonary fibrosis since they are more susceptible to misinterpretation of shrinkage and to noise and motion artifacts. Although semi-quantitative scores are widely accepted in clinical settings, automated reading might offer advancements in the future, though with its own set of challenges.

### Perspectives and Significance

Collectively, longitudinal micro-CT imaging facilitates noninvasive assessment of pulmonary fibrosis of the entire lung *in vivo* and serves as a valuable quantitative tool for future studies, especially to elucidate disease mechanisms and risk factors of disease progression or for preclinical evaluation of novel therapeutic strategies.

## DATA AVAILABILITY

The data presented in this study are available on request from the corresponding author.

## ACKNOWLEDGMENTS

We thank J. Schatterny, S. Butz and M. Finke (University of Heidelberg) for expert technical assistance.

## AUTHOR CONTRIBUTIONS

Conception and design: D.H.W.L., P.K., W.W., M.M., C.B., T.S., W.S., H-U.K., M.A.M., J.D. and M.O.W. Acquisition, analysis and interpretation of data: D.H.W.L., P.K., W.W., M.M., C.B., T.S., W.S., H-U.K., M.A.M., J.D. and M.O.W. Writing the manuscript or revising it critically for important intellectual content: D.H.W.L., P.K., W.W., M.M., C.B., T.S., W.S., H-U.K., M.A.M., J.D. and M.O.W.

