## Supplemental Table S1 for "Longitudinal Micro-Computed Tomography Detects Onset and Progression of Pulmonary Fibrosis in Conditional *Nedd4-2* Deficient Mice"

| | $\kappa$ |
|---|---|
| ***In vivo* micro-CT** | |
| Number of consolidations | 0.83 |
| Area of consolidations | 0.83 |
| **Fibrosis score** | 0.79 |
| ***Post mortem* micro-CT** | |
| Consolidation | 0.96 |
| Reticulations | 0.63 |
| Honeycombing | 0.84 |
| Peripheral bronchiectasis | 0.76 |
| Pleural lines | 0.65 |
| Fissural thickening | 0.86 |
| **Fibrosis score** | 0.40 |

**Supplemental Table S1. Interobserver agreement.** Interobserver agreement was calculated with using the weighted Cohens kappa coefficients ($\kappa$).
