## Supplemental Fig. S1 for "Longitudinal Micro-Computed Tomography Detects Onset and Progression of Pulmonary Fibrosis in Conditional *Nedd4-2* Deficient Mice"

Supplementary Figure 1

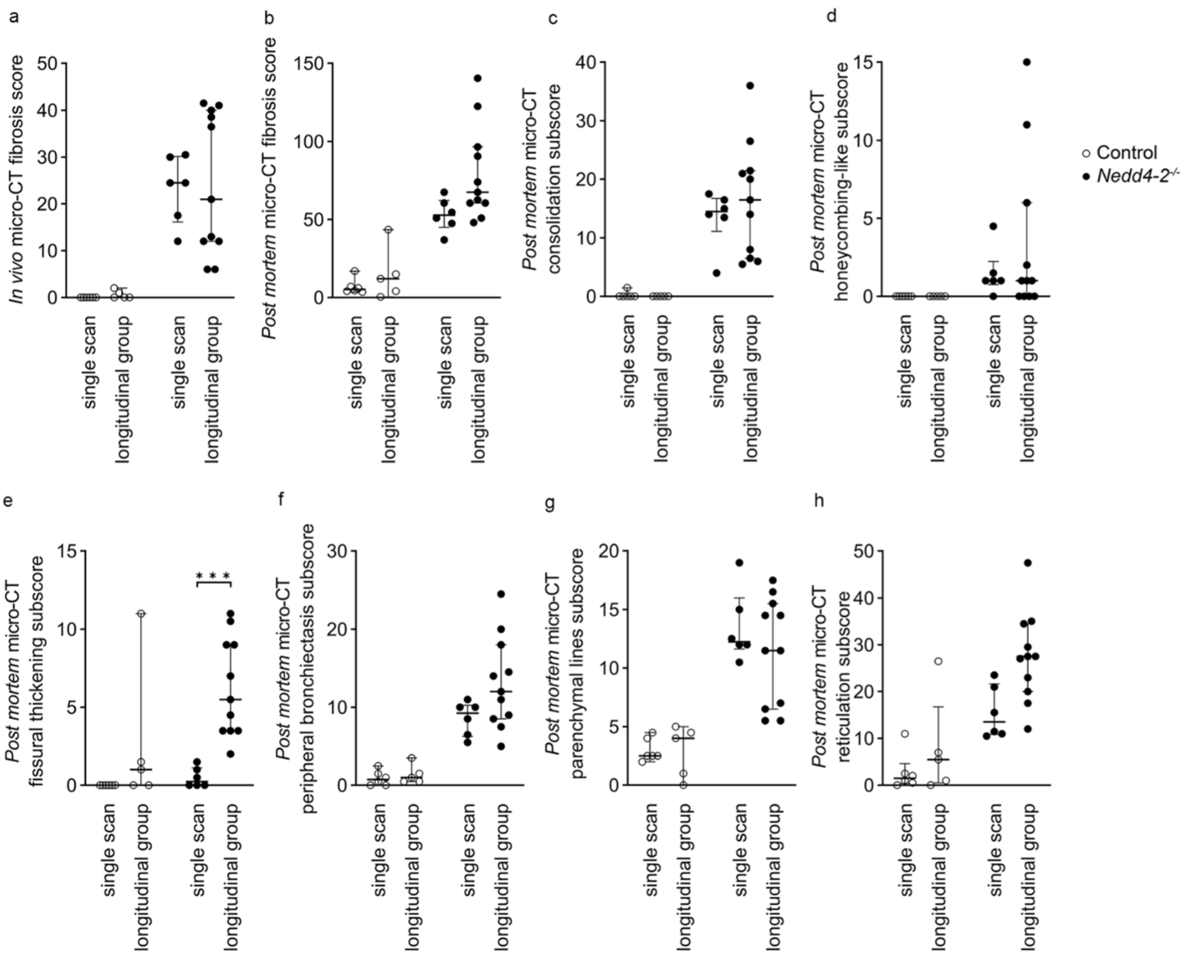


**Supplemental Fig. S1. Assessment of radiation-induced lung damages after serial *in vivo* micro-CT imaging.** (a) *In vivo* and (b) *post mortem* micro-CT fibrosis scores as well as corresponding subscores determing consolidation (c), honeycombing-like lesions (d), fissural thickening (e), peripheral bronchiectasis (f), parenchymal lines (g) and reticulations (h) in 5 months doxycycline induced conditional *Nedd4-2*$^{-/-}$ and control mice after single or repeated *in vivo* micro-CT imaging. n = 5-11 mice/group. *$^{*}P < 0.05$, $^{***}P < 0.001$.*
